# Neural network-based predictions of antimicrobial resistance in *Salmonella* spp. using k-mers counting from whole-genome sequences

**DOI:** 10.1101/2021.08.10.455825

**Authors:** Cristian C. Barros

**Affiliations:** Instituto de Ciências Biomédicas, Universidade de São Paulo, CEP 05508-000 São Paulo, Brazil

**Keywords:** Antimicrobial Resistance, Minimum Inhibitory Concentration, Artificial Neural Networks, *Salmonella*

## Abstract

Artificial intelligence-based predictions have emerged as a friendly and reliable tool for the surveillance of the antimicrobial resistance (AMR) worldwide. In this regard, genome databases typically include whole-genome sequencing (WGS) data containing AMR meta-data that can be used to train machine learning (ML) models, in order to predict phenotype features from genome samples. In this study, using a Neural Network (NN) architecture and the SGD-ADAM algorithm, we build ML antibiotic resistance models that can predict Minimum Inhibitory Concentrations (MICs) and antimicrobial susceptibility profiles of *Salmonella* spp. Data analysis was based on 7,268 genomes publicly available in PATRIC database, containing about 75,000 AMR annotations. ML models were built using reference-free *k*-mer analysis of whole-genome sequences, MIC measurements and susceptibility categories, obtaining robust and accurate results for 9 antibiotics belonging to beta-lactam, fluoroquinolone, phenicol, aminoglycoside, tetracycline and sulphonamide classes. Although the accuracy of predicting the actual MIC reaches modest levels, the within ± 1 2-fold dilution accuracy per antibiotic reaches significant levels with values that varies from 85% to 95%, with narrow 95% CIs of about 5% and individual accuracies per MIC ≳ 80%. For differentiation between “susceptible” and “resistant” values, by measuring the accuracy and error of model’s susceptibility predictions to different antibiotics, the accuracy is the same as before and ranges from 85% to 95%, with 95% CIs of about 5%, the recall extends from 75% to 85%, the precision from 60% to 90%, whereas the very major error is ≲ 20%. In summary, these results show that NN-based models are able to learn and predict the AMR phenotype from bacterial genomes based on a gene-free *k*-mer analysis.

## Introduction

Nowadays, the rapid increase of antimicrobial resistance (AMR) in clinically significant bacterial pathogens has become a public health issue. In the front line against AMR, clinical microbiology laboratories monitor the occurrence and spread of antimicrobial-resistant pathogens causing infectious diseases, and perform antimicrobial susceptibility tests (ASTs) to accurately determine which specific antibiotic an infective agent is sensitive to. The clinical testing requires the isolation, inoculation and incubation of the infective specimen to perform taxonomic identification, and susceptibility determination, including determination of the minimum inhibitory concentration (MIC). This may take from few to several days before starting to treat patients potentially infected by resistant pathogens. In this context, the construction of machine learning (ML) models that could help speed up AST results with reliable predictions has become more relevant.

Among the recently most used methods for AST predictions are those that take advantage of WGS data and ML algorithms where two general approaches are distinguished: the so-called reference-based approach, based on a set of priory known conferring resistance genes, and the reference-free approach, based on feature selection by statistical analysis. Although both approaches may work similarly, the former has the advantage of being a lightweight analysis, requiring less time and computational resources for implementation; whereas the latter is more expensive, but it could help to identify known resistant genes, as well as novel intergenomic regions associated with acquired resistance.

Recent gene-based studies focused on predicting either the AMR phenotype or the antibiotic MIC for the most critical bacteria, such as *Mycobacterium tuberculosis, Enterobacterales* (i.e., *Klebsiella pneumoniae, Escherichia coli* and *Salmonella* spp.), non-fermentative bacteria, *Staphylococcus aureus, Neisseria gonorrhoeae*, and *Streptococcus pneumoniae* [1–7]. On the other hand, gene-free studies include MIC prediction for *K. pneumoniae* to 20 antibiotics [8], MIC prediction for *Nontyphoidal Salmonella* to 15 antibiotics [9], and the PATRIC studies describing ML classifiers to 9, 13 and 16 antibiotics [10–12].

## Methods

In this study, we build AMR predictors based on a gene-free approach, since we understand that this can cover the referenced-based approach when the set of selected best features within the genome is large enough to include all those features from the intergenic region. In particular, we aim to build ML classifiers to predict MICs and antibiotic susceptibility profiles of sequenced strains. The analysis was based on 7,268 *Salmonella* genomes and near 75,000 AMR annotations, publicly available on PATRIC database (https://www.patricbrc.org/). The Salmonella collection is the most abundant collection of AMR metadata at PATRIC and uncovers a representative number of antibiotic-resistant strains. This data has been also the basis for the machine learning techniques implemented by PATRIC, focusing on antimicrobial resistance [13].

To build our models we perform a reference-free *k*-mer analysis of whole-genome sequences and consider the nucleotide *k*-mer counts and the AMR metadata as training features of the model. Although MIC data available in PATRIC covers a broad range of MIC measurements, it is an imbalanced dataset and disconnected in term of serial dilutions. Thus, in order to combine data from different sources and estimate the model quality within three two-fold dilutions, we approach those values to those of a two-fold dilution series. Determining the accuracy within ± 1 2-fold dilution is consistent with the observation that 90% to 95% of MIC results are ± 1 dilution from the median or mode for most antimicrobial/organism combinations. This is a commonly used criteria for evaluating AST systems and consistent with FDA recommendations [8, 9, 14, 15]. Here, we also include measures of the actual accuracy or raw accuracy ^1^, the multiclass ROC curve, the multiclass ROC-AUC and the multiclass F1-score.

In PATRIC, the AMR phenotype of bacterial strains is reported as being “susceptible”, “intermediate” or “resistant” to a specific antibiotic. These rates are susceptibility categories informed by researchers based on MIC data and additional clinical information. Since these interpretations depend on more variables, it is usual to find differences in the informed AMR phenotype of strains that have the same MIC. Here, in order to further classify an infection as “susceptible” or “resistant”, we consider CLSI MIC breakpoints [16] to remove the susceptibility overlap and to include also those strains that only have MIC annotations. However, we consider some empirical MIC thresholds in order to alleviate the tension between CLSI thresholds and about 2,500 strains reported as “resistant” with a lower MIC value. Furthermore, we consider strains of intermediate resistance as “resistant”. Although the model is build in the basis of MIC measurements, we translate those measurements to susceptibility categories using MIC breakpoints and estimate the model quality by measuring the accuracy, recall, precision, F1-score, major error (ME) and very major error (VME) of model’s binary predictions. We include results to 9 antibiotics for which the model is effectively able to learn to identify both susceptible and resistant strains. These include 4 beta-lactams (*i*.*e*., ceftriaxone, ampicillin, amoxicillin/clavulanic acid ^2^ and cefoxitin) and 5 from other families (*i*.*e*., criprofloxacin, chloramphenicol, gentamicin, tetracycline and sulfisoxazole).

This work is divided as follows: In Section 1 we describe the genome data and AMR metadata available in PATRIC and define MIC breakpoints to support susceptibility predictions. In Section 2 we describe our AMR classifiers arguing about the ML algorithm, the *k*-mer analysis and feature selection, as well as the metrics used to evaluate the model’s performance. Then, in Section 3, we present the results obtained for such metrics as average measurements of a 10-fold cross-validation. Finally, in Section 4, we discuss the validity of the model.

### 1 AMR metadata

AMR metadata at PATRIC is in the form of MICs or SIR terms. The MIC is defined as the minimum concentration of an antibiotic to prevent the further growth of an infectious agent *in vitro*. MIC measurements are expressed in units of mg/L, and here, we approximate their values to those of a 2-fold dilution series. This will reduce the MIC classes per antibiotic and simplify the output of machine learning models. SIR terms represent the degree of resistance of infectious agents under the action of antibiotics. These are critical reports made by researchers based on clinical information and established for at least four international committees/organizations, ^3^, in order to simplify and standardize the categorization of infectious agents all over the world. S, I and R classifications, as stated in the ISO norms, 20776-1 and 20776-2 [17], and in the CLSI documents [16], are defined as follows:

- Susceptible (S): A bacterial strain is said to be susceptible to a given antibiotic when it is inhibited *in vitro* by a concentration of this drug that is associated with a high likelihood of therapeutic success.
- Intermediate (I): The sensitivity of a bacterial strain to a given antibiotic is said to be intermediate when it is inhibited *in vitro* by a concentration of this drug that is associated with an uncertain therapeutic effect.
- Resistant (R): A bacterial strain is said to be resistant to a given antibiotic when it is inhibited *in vitro* by a concentration of this drug that is associated with a high likelihood of therapeutic failure.

#### 1.1 MIC breakpoints

MIC breakpoints, as described in the CLSI documents for AST standards [16], are the MIC values used to determine whether an organism is “susceptible”, “intermediate” or “resistant” to a specific antibiotic. ^4^ For “susceptible” isolates, the MIC is smaller than the lower threshold and likely to be effective in treatments more than 90% of the time. For “intermediate” isolates, therapeutic treatments are effective at higher doses or at the lower threshold if antimicrobial concentrates at tissue site. For “resistant” bacteria, the MIC values are greater than a higher threshold and unlikely to be able to achieve effective levels of the drug at safe doses.

The susceptibility labels attached to most lab reports are based on MIC breakpoints and are characteristics of particular species to specific antibiotics. However, much more AMR metadata corresponds to MIC measurements than to Susceptibility categories. SIR terms are general definitions to monitor antibiotic resistance and robust indicator of clinical success or failure, therefore *in vitro* MIC measurements are not suffice to classify an isolated strain as S, I or R. The right classification must consider all clinical and pharmacological information available, which includes infection indications, dosage, pharmacokinetics and pharmacodynamics. Nevertheless, we can relate directly S, I or R terms to MIC measurements by using established CLSI MIC breakpoints. The aim is to include all data samples with associated MIC measurements and to eliminate the overlap between reported susceptibility rates.

However, in this analysis we adopt a combination of empirical and established MIC breakpoints. For the lower thresholds, we consider MIC breakpoints from CLSI standards to all antibiotics but sulfisoxazole to which we use a lower threshold by one two-fold dilution. For the higher thresholds we use the CLSI MIC breakpoints to 7 antibiotics and empirical breakpoints for criprofloxacin and sulfisoxazole to which we find lower thresholds. For ciprofloxacin we find 476 genomes rated as “resistent” with a MIC value lower by 1 to 2 two-fold dilutions than the established breakpoint, while for sulfisoxazole we find 2,102 genomes reported as “resistent” with a MIC value lower by 1 two-fold dilution than the established breakpoint. Thus, we adopt the MIC breakpoints showed in Table 1.

**Table 1:** MIC breakpoints and susceptibility rating. For the lower thresholds, we consider MIC breakpoints from CLSI standards to all antibiotics but sulfisoxazole to which we use a MIC value of 128 mg/L instead of 256 mg/L. For the higher thresholds we use CLSI breakpoints for 7 out of 9 antibiotics. For ciprofloxacin we use the MIC value of 0.25 mg/L instead of 1 mg/L, whereas for sulfisoxazole we use 256 mg/L instead of 512 mg/L.

| Antibiotic | *Salmonella* ratings (MIC in mg/L) |  |  |
| --- | --- | --- | --- |
|  | Susceptible | Intermediate | Resistant |
| 1 ciprofloxacin | $\leq 0.06$ | 0.125 | $\geq 0.25$ |
| 2 ceftriaxone | $\leq 1$ | 2 | $\geq 4$ |
| 3 gentamicin | $\leq 4$ | 8 | $\geq 16$ |
| 4 ampicillin | $\leq 8$ | 16 | $\geq 32$ |
| 5 amoxicillin/clavulanic acid | $\leq 8$ | 16 | $\geq 32$ |
| 6 cefoxitin | $\leq 8$ | 16 | $\geq 32$ |
| 7 tetracycline | $\leq 4$ | 8 | $\geq 16$ |
| 8 chloramphenicol | $\leq 8$ | 16 | $\geq 32$ |
| 9 sulfisoxazole | $\leq 128$ | – | $\geq 256$ |

Applying this strategy to those antibiotics that present more than 100 strains per MIC, we obtain the MIC distribution and susceptibility map showed in Fig. 1. This map shows the number of genomes available per antibiotic and MIC and the corresponding susceptibility rating.

**Figure 1:**
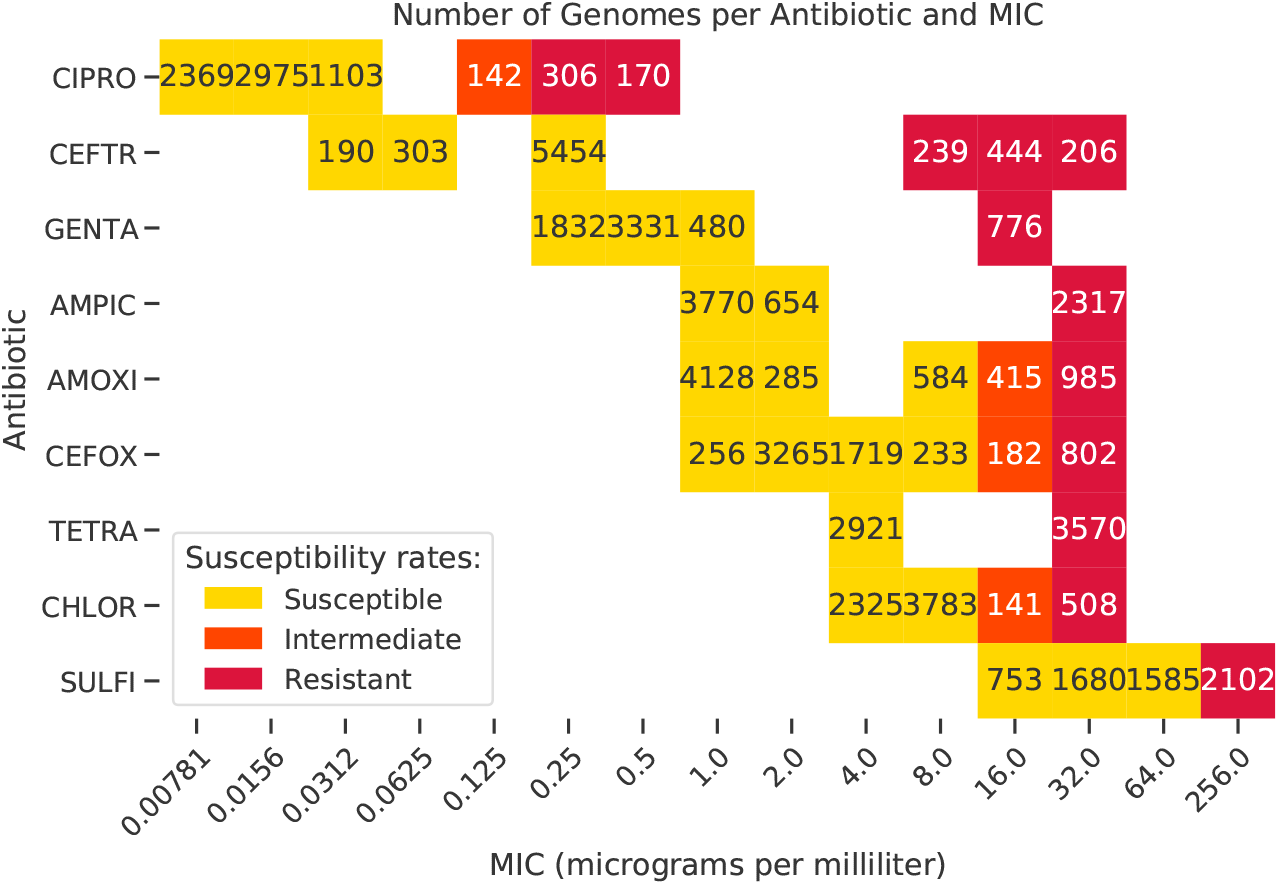
MIC and susceptibility distribution. Number of genomes available per antibiotic and MIC, and per susceptibility categories, according to the MIC breakpoints in Table 1. We only retain data for those MIC classes that have more than 100 samples available.

### 2 AMR classifiers

AMR classifiers for *Salmonella* species to 9 antibiotics were constructed based on an artificial Neural Network, implemented in Keras, and on the SGD-ADAM algorithm [18]. The network architecture is of single-layer neural network of 64 neurons width. We consider the *k*-mers counts of genome assemblies as the input of the model, and the corresponding antibiotic MIC measurements as the output. Here, *k*-mers means subsequences of length *k* contained in a genome assembly, in which *k*-mers are composed of nucleotides (*i*.*e*., A, T, G, and C). The number of *k*-mers in a biological sequence of length *L* is *L* − *k* + 1, being the number of unique *k*-mers less than the total number in a sample from repetitions throughout the sequence. These repetitions are precisely the counts carrying the information provided to the model. The value of *k* is representative of how deep and refined the genomic analysis is. For this analysis we consider *k* = 10. This is commonly used starting value for a high performance. The *k*-mers counts was produced using only Python routines and Unix shell scripts. A developed software performs the *k*-mer analysis, the statistical analysis for feature selection, model building and K-fold cross-validation. This allows the pre-processing of thousands of genomes in regular desk computers, transforming the genome data into spectral data. This will be available soon on GitHub.

When building an ML model we start by defining the validation procedure with which to measure the model performance. Here we will evaluate the model using a 10-fold cross validation. For this purpose, the entire dataset is divided into 10 folds, and used alternately as training, validation and test datasets. At each fold round, 8 folds are used to train or fit the network, 1 fold to evaluate the performance during training and 1 fold is used to evaluate the model on unseen data. Thus, we estimate the model performance obtaining multiple measurements of model quality.

Models were constructed in the gene-free approach where the most relevant nucleotide *k*-mers are selected from statistical analysis. The initial set of *k*-mers counts are arranged as rows into a matrix, and for each given antibiotic, the number of columns is reduced by a statistical F-test analysis. The model accuracy strongly depends on the number of selected *k*-mers, reaching a high performance for values 500.

#### 2.1 Accuracy

In this work, we evaluate the model’s performance to all antibiotics by a 10-fold cross-validation experiment, measuring the average of statistical precision and error metrics and their 95% CIs.

For the model’s MIC predictions we evaluate the multi-class Receiver Operating Characteristic (ROC) curves, the raw accuracy, the accuracy within ±1 two-fold dilution, the F1-score and the area under the ROC curves (ROC-AUCs). The raw accuracy is defined as the accuracy of predicting the actual MIC, whereas the accuracy within ± 1 2-fold dilution is the accuracy of predicting either the actual MIC, the successor MIC or the predecessor MIC, in a 2-fold dilution series. An ROC curve illustrates the performance ability plotting the True Positive Rate (TPR) versus the False Positive Rate (FPR), where a functional model presents the TPR growing faster than the FPR up to a particular value of the logistic regression function – the classification threshold. This curve is commonly defined for binary problems, however, it can be expanded to multiclass problems considering as positive one class versus all at a time. The ROC curve plots the TPR in the *y* axis and the FPR in the *x* axis (Figs. 4 and 5). This means that the top left corner of the plot is the “ideal” point – a FPR of zero, and a TPR of one. This point defines the ideal area under the curve that is equal to 1. Thus, the ROC-AUC is also a measure of performance, measuring the quality across all possible classification thresholds. Together with the ROC-AUC we include the multi-class F1-score, which is the harmonic mean of Precision and Recall extended to the multi-class case.

To measure the quality of model’s binary predictions, we simply translate the true and predicted MIC values to susceptibility categories, using the MIC breakpoints, and considering strains of intermediate resistance as “resistant”. Then, considering the “resistant” label as the positive class, we compute the accuracy, recall, precision, F1-score, major error (ME) and very major error (VME) of susceptibility predictions. The accuracy is the weighted average of the particular accuracies of predicting “susceptible” and “resistant” strains, the recall is the accuracy of predicting “resistant” strains, whereas the precision corresponds to the percentage of predicted resistant strains correctly classified. While recall stands for the sensitivity of the model, precision stands for the probability that a particular strain identified as resistant is actually resistant. Concerning error metrics, the ME is defined as the percentage of true susceptible strains predicted as resistant and the VME the percentage of true resistant strains predicted as susceptible.

### 3 Results

Results for the metrics described above are presented as average measurements of a 10-fold cross validation. In Fig. 2 the within ± 1 2-fold dilution accuracy per antibiotic and MIC is showed. In this regard, the mean accuracy per antibiotic varies from 85% to 95%, with narrow 95% CIs of about 5% (Table 4) and individual accuracies per MIC ≳ 80% (Fig. 2). The highest and most balanced results are found for Tetracycline, reaching exact per MIC accuracies of 89% and 88% for MICs equal to 4 *µ*g/mL (S) and 32 *µ*g/mL (R), respectively. Measurements of this metric can be compared to the raw accuracy in Table 2. Although the raw accuracy reaches modest values, all values are over chance but the combinations co-amoxiclav/2, cefoxitin/8 chloramphenicol/16, which have the least amount of samples per MIC. Figs. 4 and 5 show the multiclass ROC curves to all antibiotic models, as well as the ROC-AUC- and F1-scores. In this respect, we can see that all models are functional with considerable ROC-AUC scores and fast growing TPRs – up to a certain value of the classification threshold. The F1-score and ROC-AUC are also summarized in Table 3.

**Figure 2:**
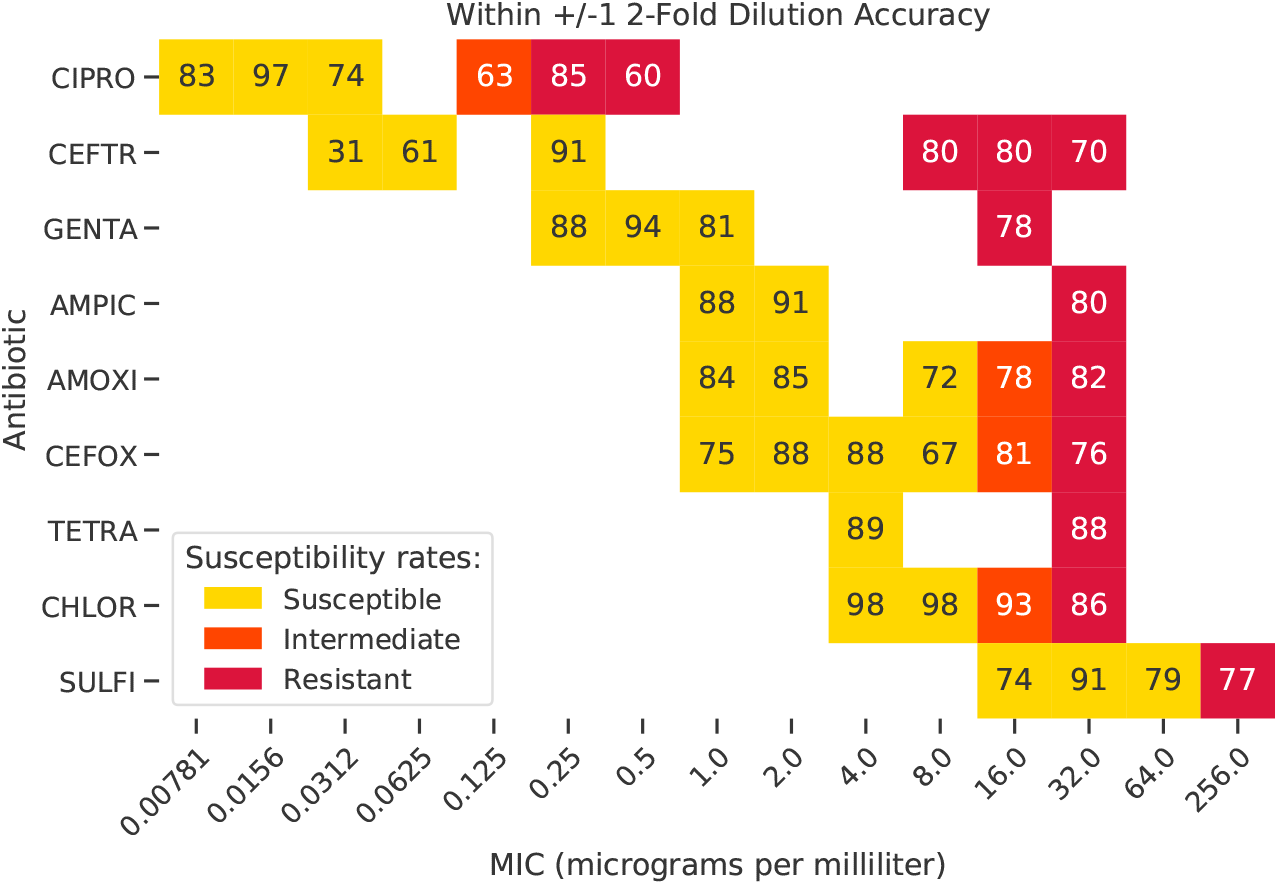
The within ± 1 2-fold dilution accuracy obtained to 9 antibiotics. Significantly, the accuracy obtained for MIC classes rated as “resistant” are greater than 80%.

**Table 2:** Raw accuracy and within ± 1 2-fold dilution accuracy per antibiotic and MIC class as average measurement of a 10-fold cross-validation experiment, together with their 95% CIs and the number of samples available per class.

| Antibiotic | MIC | Genome samples | Raw Accuracy | Raw Accuracy 95% CI | Within $\pm 1$ Two-fold Accuracy | Within $\pm 1$ Two-fold accuracy 95% CI |
| --- | --- | --- | --- | --- | --- | --- |
| ciprofloxacin | 0.00781 | 2369 | 0.65 | [0.59, 0.72] | 0.83 | [0.78, 0.89] |
| ciprofloxacin | 0.0156 | 2975 | 0.6 | [0.53, 0.67] | 0.97 | [0.95, 0.99] |
| ciprofloxacin | 0.0312 | 1103 | 0.51 | [0.46, 0.57] | 0.74 | [0.70, 0.78] |
| ciprofloxacin | 0.125 | 142 | 0.21 | [0.09, 0.34] | 0.63 | [0.41, 0.86] |
| ciprofloxacin | 0.25 | 306 | 0.46 | [0.30, 0.63] | 0.85 | [0.78, 0.91] |
| ciprofloxacin | 0.5 | 170 | 0.24 | [0.11, 0.37] | 0.6 | [0.21, 0.99] |
| ceftriaxone | 0.0312 | 190 | 0.31 | [0.22, 0.41] | 0.31 | [0.22, 0.41] |
| ceftriaxone | 0.0625 | 303 | 0.61 | [0.42, 0.80] | 0.61 | [0.42, 0.80] |
| ceftriaxone | 0.25 | 5454 | 0.91 | [0.88, 0.94] | 0.91 | [0.88, 0.94] |
| ceftriaxone | 8 | 239 | 0.59 | [0.35, 0.83] | 0.8 | [0.63, 0.98] |
| ceftriaxone | 16 | 444 | 0.49 | [0.40, 0.59] | 0.8 | [0.70, 0.90] |
| ceftriaxone | 32 | 206 | 0.24 | [-0.03, 0.51] | 0.7 | [0.42, 0.97] |
| gentamicin | 0.25 | 1832 | 0.55 | [0.48, 0.61] | 0.88 | [0.83, 0.94] |
| gentamicin | 0.5 | 3331 | 0.5 | [0.43, 0.57] | 0.94 | [0.92, 0.97] |
| gentamicin | 1 | 480 | 0.31 | [0.19, 0.43] | 0.81 | [0.64, 0.97] |
| gentamicin | 16 | 776 | 0.78 | [0.70, 0.87] | 0.78 | [0.70, 0.87] |
| ampicillin | 1 | 3770 | 0.71 | [0.67, 0.76] | 0.88 | [0.86, 0.90] |
| ampicillin | 2 | 654 | 0.48 | [0.32, 0.63] | 0.91 | [0.84, 0.97] |
| ampicillin | 32 | 2317 | 0.8 | [0.74, 0.86] | 0.8 | [0.74, 0.86] |
| co-amoxiclav | 1 | 4128 | 0.74 | [0.71, 0.78] | 0.84 | [0.81, 0.87] |
| co-amoxiclav | 2 | 285 | 0.18 | [0.04, 0.33] | 0.85 | [0.73, 0.98] |
| co-amoxiclav | 8 | 584 | 0.61 | [0.51, 0.71] | 0.72 | [0.63, 0.82] |
| co-amoxiclav | 16 | 415 | 0.53 | [0.39, 0.67] | 0.78 | [0.65, 0.91] |
| co-amoxiclav | 32 | 985 | 0.79 | [0.69, 0.88] | 0.82 | [0.75, 0.88] |
| cefoxitin | 1 | 256 | 0.27 | [0.08, 0.46] | 0.75 | [0.58, 0.91] |
| cefoxitin | 2 | 3265 | 0.64 | [0.60, 0.68] | 0.88 | [0.85, 0.90] |
| cefoxitin | 4 | 1719 | 0.53 | [0.46, 0.60] | 0.88 | [0.83, 0.93] |
| cefoxitin | 8 | 233 | 0.14 | [0.02, 0.26] | 0.67 | [0.49, 0.84] |
| cefoxitin | 16 | 182 | 0.27 | [0.12, 0.41] | 0.81 | [0.66, 0.95] |
| cefoxitin | 32 | 802 | 0.63 | [0.53, 0.73] | 0.76 | [0.67, 0.85] |
| tetracycline | 4 | 2921 | 0.89 | [0.85, 0.93] | 0.89 | [0.85, 0.93] |
| tetracycline | 32 | 3570 | 0.88 | [0.83, 0.92] | 0.88 | [0.83, 0.92] |
| chloramphenicol | 4 | 2325 | 0.67 | [0.63, 0.72] | 0.98 | [0.97, 0.99] |
| chloramphenicol | 8 | 3783 | 0.74 | [0.69, 0.79] | 0.98 | [0.97, 0.99] |
| chloramphenicol | 16 | 141 | 0.12 | [-0.03, 0.28] | 0.93 | [0.78, 1.09] |
| chloramphenicol | 32 | 508 | 0.8 | [0.70, 0.90] | 0.86 | [0.80, 0.91] |
| sulfisoxazole | 16 | 753 | 0.45 | [0.29, 0.61] | 0.74 | [0.62, 0.86] |
| sulfisoxazole | 32 | 1680 | 0.36 | [0.28, 0.44] | 0.91 | [0.87, 0.96] |
| sulfisoxazole | 64 | 1585 | 0.49 | [0.41, 0.58] | 0.79 | [0.72, 0.86] |
| sulfisoxazole | 256 | 2102 | 0.77 | [0.70, 0.84] | 0.77 | [0.70, 0.84] |

**Table 3:** Multiclass F1-score and ROC-AUC as average measurement of a 10-fold cross-validation experiment, together with their 95% CIs and the number of samples available per class.

| Antibiotic | MIC | Genome samples | F1-score | F1-score 95% CI | ROC-AUC | ROC-AUC 95% CI |
| --- | --- | --- | --- | --- | --- | --- |
| ciprofloxacin | 0.00781 | 2369 | 0.64 | [0.57, 0.70] | 0.82 | [0.79, 0.86] |
| ciprofloxacin | 0.0156 | 2975 | 0.65 | [0.60, 0.71] | 0.79 | [0.77, 0.82] |
| ciprofloxacin | 0.0312 | 1103 | 0.45 | [0.40, 0.50] | 0.77 | [0.72, 0.81] |
| ciprofloxacin | 0.125 | 142 | 0.21 | [0.03, 0.39] | 0.88 | [0.79, 0.98] |
| ciprofloxacin | 0.25 | 306 | 0.45 | [0.31, 0.58] | 0.92 | [0.87, 0.98] |
| ciprofloxacin | 0.5 | 170 | 0.23 | [0.09, 0.37] | 0.87 | [0.74, 1.01] |
| ceftriaxone | 0.0312 | 190 | 0.33 | [0.20, 0.45] | 0.96 | [0.93, 1.00] |
| ceftriaxone | 0.0625 | 303 | 0.59 | [0.44, 0.75] | 0.98 | [0.98, 0.99] |
| ceftriaxone | 0.25 | 5454 | 0.94 | [0.92, 0.95] | 0.96 | [0.95, 0.97] |
| ceftriaxone | 8 | 239 | 0.44 | [0.30, 0.59] | 0.9 | [0.83, 0.97] |
| ceftriaxone | 16 | 444 | 0.42 | [0.32, 0.53] | 0.88 | [0.85, 0.92] |
| ceftriaxone | 32 | 206 | 0.23 | [-0.04, 0.49] | 0.85 | [0.79, 0.91] |
| gentamicin | 0.25 | 1832 | 0.52 | [0.47, 0.57] | 0.74 | [0.70, 0.77] |
| gentamicin | 0.5 | 3331 | 0.56 | [0.52, 0.61] | 0.66 | [0.62, 0.70] |
| gentamicin | 1 | 480 | 0.22 | [0.12, 0.32] | 0.69 | [0.59, 0.79] |
| gentamicin | 16 | 776 | 0.72 | [0.64, 0.80] | 0.94 | [0.90, 0.97] |
| ampicillin | 1 | 3770 | 0.76 | [0.72, 0.80] | 0.83 | [0.81, 0.86] |
| ampicillin | 2 | 654 | 0.36 | [0.26, 0.46] | 0.76 | [0.73, 0.80] |
| ampicillin | 32 | 2317 | 0.79 | [0.76, 0.82] | 0.92 | [0.90, 0.94] |
| co-amoxiclav | 1 | 4128 | 0.8 | [0.77, 0.83] | 0.84 | [0.81, 0.88] |
| co-amoxiclav | 2 | 285 | 0.13 | [0.03, 0.24] | 0.66 | [0.56, 0.76] |
| co-amoxiclav | 8 | 584 | 0.55 | [0.48, 0.63] | 0.87 | [0.83, 0.91] |
| co-amoxiclav | 16 | 415 | 0.44 | [0.31, 0.57] | 0.87 | [0.82, 0.92] |
| co-amoxiclav | 32 | 985 | 0.75 | [0.69, 0.82] | 0.94 | [0.92, 0.97] |
| cefoxitin | 1 | 256 | 0.18 | [0.06, 0.29] | 0.77 | [0.68, 0.87] |
| cefoxitin | 2 | 3265 | 0.68 | [0.66, 0.71] | 0.79 | [0.77, 0.82] |
| cefoxitin | 4 | 1719 | 0.55 | [0.50, 0.61] | 0.78 | [0.75, 0.81] |
| cefoxitin | 8 | 233 | 0.12 | [0.01, 0.22] | 0.68 | [0.58, 0.79] |
| cefoxitin | 16 | 182 | 0.2 | [0.09, 0.31] | 0.8 | [0.70, 0.90] |
| cefoxitin | 32 | 802 | 0.58 | [0.52, 0.65] | 0.9 | [0.87, 0.94] |
| tetracycline | 4 | 2921 | 0.87 | [0.84, 0.91] | 0.95 | [0.93, 0.97] |
| tetracycline | 32 | 3570 | 0.89 | [0.86, 0.92] | 0.95 | [0.93, 0.97] |
| chloramphenicol | 4 | 2325 | 0.66 | [0.62, 0.71] | 0.83 | [0.81, 0.85] |
| chloramphenicol | 8 | 3783 | 0.75 | [0.73, 0.77] | 0.8 | [0.78, 0.82] |
| chloramphenicol | 16 | 141 | 0.11 | [-0.02, 0.23] | 0.73 | [0.59, 0.86] |
| chloramphenicol | 32 | 508 | 0.76 | [0.70, 0.82] | 0.96 | [0.93, 1.00] |
| sulfisoxazole | 16 | 753 | 0.37 | [0.26, 0.47] | 0.78 | [0.72, 0.83] |
| sulfisoxazole | 32 | 1680 | 0.39 | [0.32, 0.45] | 0.7 | [0.66, 0.73] |
| sulfisoxazole | 64 | 1585 | 0.48 | [0.41, 0.56] | 0.77 | [0.72, 0.81] |
| sulfisoxazole | 256 | 2102 | 0.8 | [0.75, 0.86] | 0.93 | [0.90, 0.95] |

**Figure 3:**
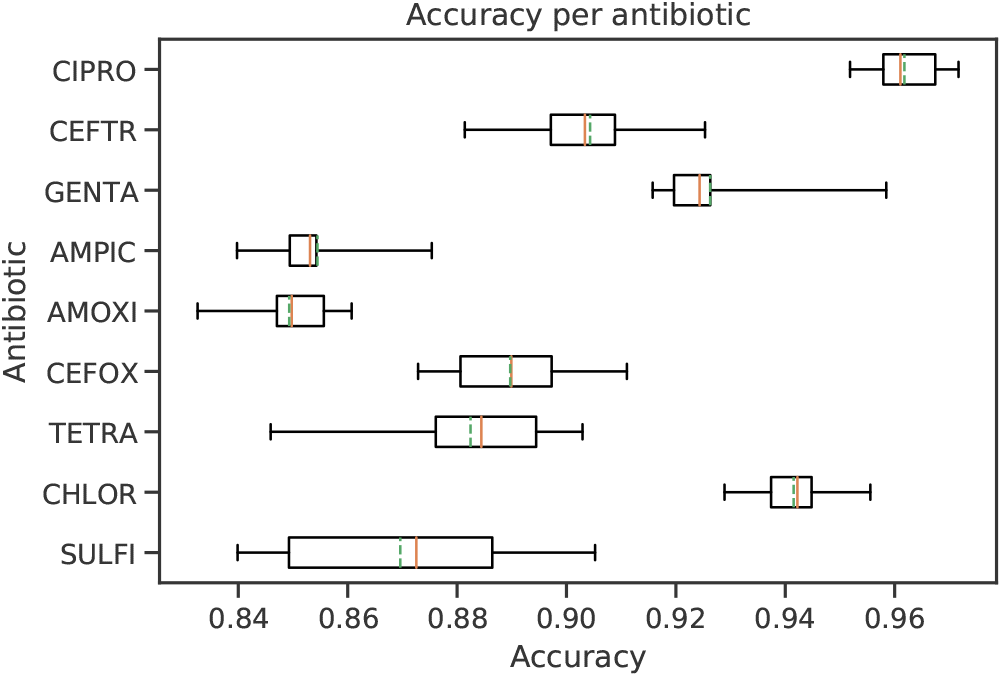
The accuracy obtained from 10-fold cross-validation to all antibiotics. The box extends from the first to third quartile values, the orange vertical line represents the median, the green dashed line the mean and the whiskers stand for the full range of accuracy measurements.

**Figure 4:**
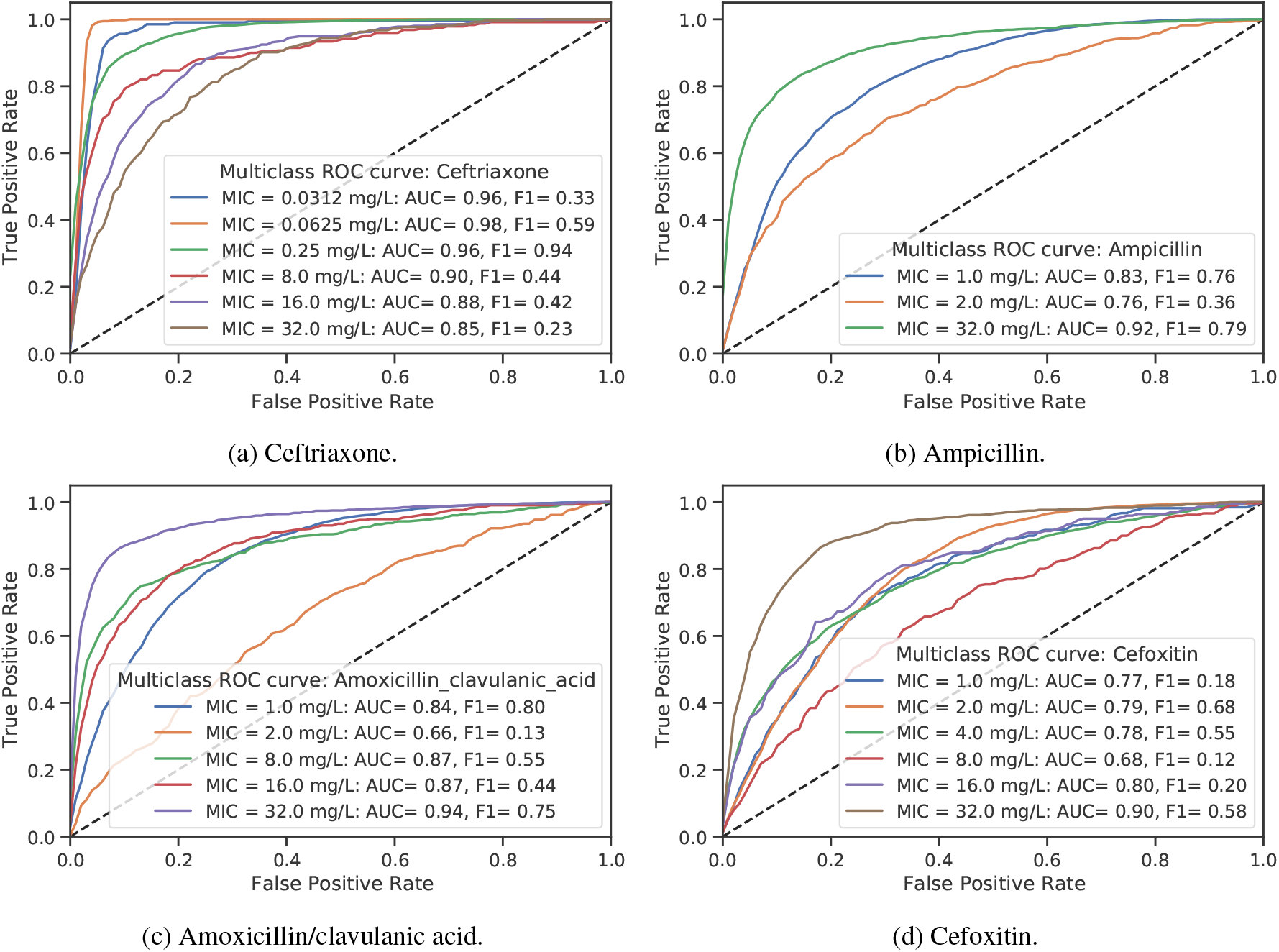
Multiclass ROC curves, ROC-AUC and F1 scores for the AMR classifiers of four beta-lactams; Ceftriaxone, Ampicillin, Amoxicillin/clavulanic acid and Cefoxitin. For all antibiotic/MIC combinations the ROC curves present considerable ROC-AUC scores and fast growing TPRs – up to a certain value of the classification threshold. The F1-scores are also shown for comparison.

**Figure 5:**
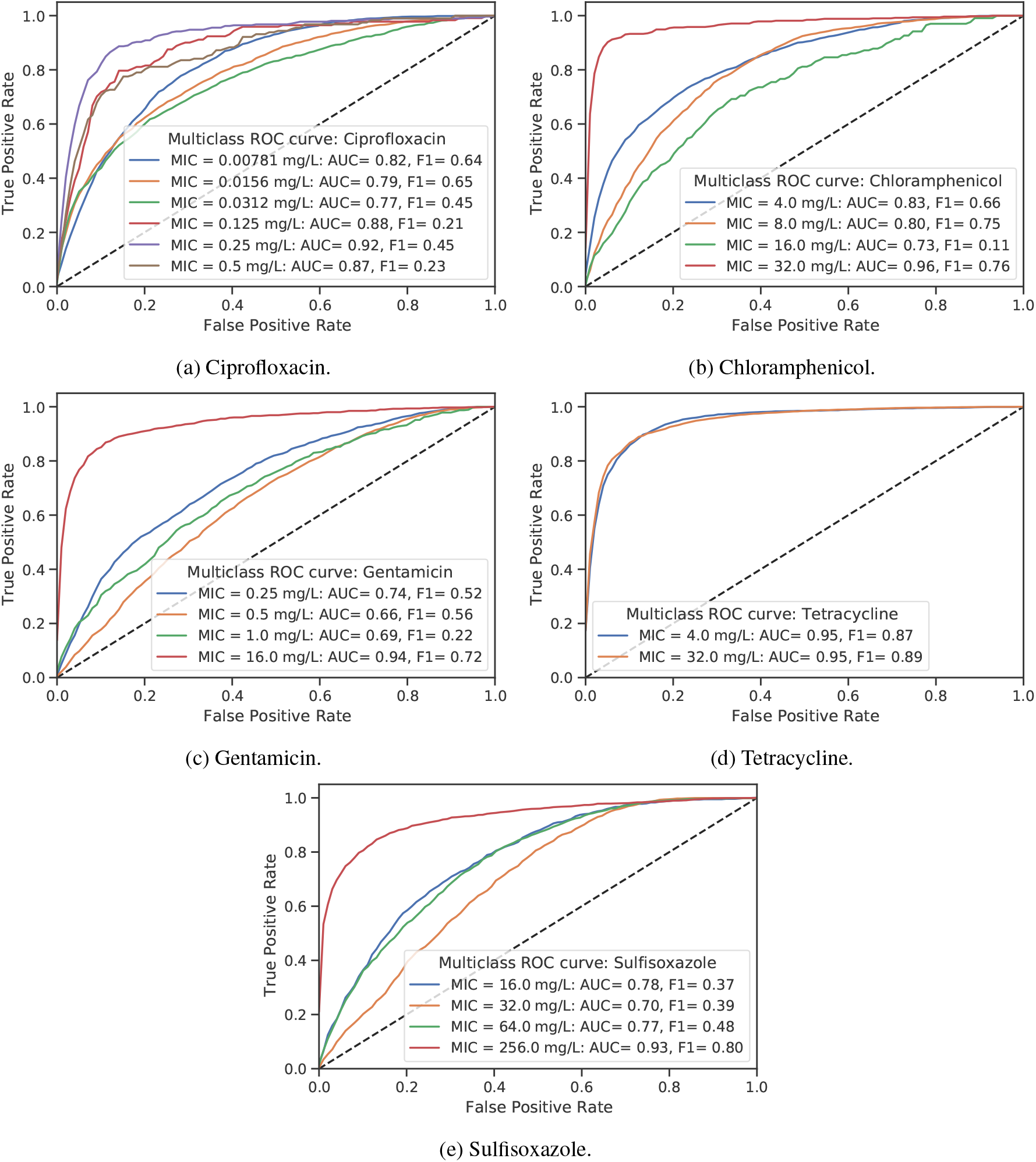
Same as Fig. 4 but for Ciprofloxacin, Chloramphenicol, Gentamicin, Tetracycline and Sulfisoxazole.

Concerning the model’s ability of predicting between susceptibility categories, Fig. 3 shows the full range of accuracy measurements from 10-fold cross-validation to all antibiotics. The box extends from the first to third quartile values, the orange vertical line represents the median, the green dashed line the mean and the whiskers stand for the full range of accuracy measurements. This plot shows roughly symmetric accuracy measurements throughout 10 folds for all antibiotic but gentamicin. Table 4 summarizes average values for accuracy, recall and precision metrics, Table 5 summarizes average values for the F1-score and the ROC-AUC, and Table 6 summarizes average values for ME and VME metrics. In the susceptibility basis, the accuracy per antibiotic is the same as before and ranges from 85% to 95%, with 95% CIs of about 5%, the recall extends from 75% to 85%, whereas the precision from 60% to 90%. The highest accuracies are for ciprofloxacin (96%), chloramphenicol (94%), gentamicin (93%) and ceftriaxone (90%), however, the best recall measurements are for tetracycline (88%), ciprofloxacin (82%), ceftriaxone (81%), and ampicillin (80%), whereas the highest precision is for tetracycline (90%). Consistently, the ME is less than 15% and the VME less than 20%. In this case, the lowest errors are for tetracycline, ciprofloxacin and ceftriaxone, with VME equal to 12%, 18% and 19%, respectively.

**Table 4:** Accuracy, recall and precision per antibiotic as average measurement of a 10-fold cross-validation experiment, together with their 95% CIs and the number of samples available per susceptibility category.

| Antibiotic | S | R | Accuracy | Accuracy 95% CI | Recall | Recall 95% CI | Precision | Precision 95% CI |
| --- | --- | --- | --- | --- | --- | --- | --- | --- |
| ciprofloxacin | 6447 | 618 | 0.96 | [0.95 0.97] | 0.82 | [0.73 0.91] | 0.76 | [0.64 0.88] |
| ceftriaxone | 5947 | 889 | 0.9 | [0.88 0.92] | 0.81 | [0.76 0.86] | 0.6 | [0.47 0.73] |
| gentamicin | 5643 | 776 | 0.93 | [0.91 0.95] | 0.78 | [0.7 0.86] | 0.67 | [0.55 0.79] |
| ampicillin | 4424 | 2317 | 0.85 | [0.83 0.87] | 0.8 | [0.74 0.86] | 0.78 | [0.74 0.82] |
| co-amoxiclav | 4997 | 1400 | 0.85 | [0.83 0.87] | 0.75 | [0.67 0.83] | 0.63 | [0.56 0.7 ] |
| cefoxitin | 5473 | 984 | 0.89 | [0.87 0.91] | 0.76 | [0.67 0.85] | 0.61 | [0.52 0.7 ] |
| tetracycline | 2921 | 3570 | 0.88 | [0.85 0.91] | 0.88 | [0.83 0.93] | 0.9 | [0.87 0.93] |
| chloramphenicol | 6108 | 649 | 0.94 | [0.92 0.96] | 0.76 | [0.67 0.85] | 0.67 | [0.58 0.76] |
| sulfisoxazole | 4018 | 2102 | 0.87 | [0.83 0.91] | 0.77 | [0.7 0.84] | 0.84 | [0.77 0.91] |

**Table 5:** F1-score and ROC-AUC per antibiotic as average measurement of a 10-fold cross-validation experi-ment, together with their 95% CIs and the number of samples available per susceptibility category.

| Antibiotic | S | R | F1-score | F1-score<br>95% CI | ROC-AUC | ROC-AUC<br>95% CI |
| --- | --- | --- | --- | --- | --- | --- |
| ciprofloxacin | 6447 | 618 | 0.79 | [0.7 0.88] | 0.9 | [0.85 0.95] |
| ceftriaxone | 5947 | 889 | 0.69 | [0.6 0.78] | 0.86 | [0.83 0.89] |
| gentamicin | 5643 | 776 | 0.72 | [0.64 0.8 ] | 0.86 | [0.82 0.9 ] |
| ampicillin | 4424 | 2317 | 0.79 | [0.76 0.82] | 0.84 | [0.81 0.87] |
| co-amoxiclav | 4997 | 1400 | 0.68 | [0.64 0.72] | 0.81 | [0.78 0.84] |
| cefoxitin | 5473 | 984 | 0.68 | [0.61 0.75] | 0.83 | [0.78 0.88] |
| tetracycline | 2921 | 3570 | 0.89 | [0.86 0.92] | 0.88 | [0.85 0.91] |
| chloramphenicol | 6108 | 649 | 0.71 | [0.64 0.78] | 0.86 | [0.82 0.9 ] |
| sulfisoxazole | 4018 | 2102 | 0.8 | [0.74 0.86] | 0.85 | [0.8 0.9] |

**Table 6:** Major error and very major error per antibiotic as average measurement of a 10-fold cross-validation experiment, together with their 95% CIs and the number of samples available per susceptibility category.

| Antibiotic | S | R | Major Error | Major Error<br>95% CI | Very Major<br>Error | Very Major<br>Error 95% CI |
| --- | --- | --- | --- | --- | --- | --- |
| ciprofloxacin | 6447 | 618 | 0.025 | [0.012 0.038] | 0.18 | [0.085 0.27 ] |
| ceftriaxone | 5947 | 889 | 0.081 | [0.053 0.11 ] | 0.19 | [0.14 0.24] |
| gentamicin | 5643 | 776 | 0.053 | [0.031 0.075] | 0.22 | [0.14 0.3 ] |
| ampicillin | 4424 | 2317 | 0.12 | [0.097 0.14 ] | 0.2 | [0.14 0.26] |
| co-amoxiclav | 4997 | 1400 | 0.12 | [0.099 0.14 ] | 0.25 | [0.17 0.33] |
| cefoxitin | 5473 | 984 | 0.086 | [0.063 0.11 ] | 0.24 | [0.15 0.33] |
| tetracycline | 2921 | 3570 | 0.11 | [0.071 0.15 ] | 0.12 | [0.075 0.17 ] |
| chloramphenicol | 6108 | 649 | 0.04 | [0.026 0.054] | 0.24 | [0.15 0.33] |
| sulfisoxazole | 4018 | 2102 | 0.079 | [0.042 0.12 ] | 0.23 | [0.16 0.3 ] |

### 4 Discussion

ML models could help microbiologists to identify resistant strains, assist physicians with rapid and accurate diagnostics for antibiotic administration and so reduce the misuse and overuse of antibiotic drugs.

In this study, we have used a Neural Network algorithm, genome data and AMR metadata to build antibiotic susceptibility predictors. We have analyzed all data available for *Salmonella* bacteria in PATRIC platform and selected data for those antibiotics that the models are effectively able to learn. Although the selected MIC metadata is still very unbalanced, neural networks can balance the results by introducing MIC weights, which is a way to train the model with MICs that needs more attention, preventing over-fitting and without requiring a sub-sampling. We perform a statistical F-test analysis in order to reduce the *k*-mer features and training time, although the *k*-mer counting and statistical analysis take most of the time. The model reach the maximum accuracy very fast requiring only from 200 to 500 *k*-mer features – out of a total of one million for *k* = 10. Notably, artificial neural networks can produce good-quality predictions from WGS data in the gene-free approach.

We identified four critical factors that influence the model performance: i) the number of samples per antibiotic and MIC, ii) the balance between MIC classes, iii) the number of MIC classes per antibiotic and iv) the length of the oligonucleotide *k*-mers. A high and balanced number of samples per antibiotic increases the accuracy of model’s predictions, with significant levels for *k* ≳ 10. Inversely, the quality of the model decreases as the number of classes increases.

Although the model architecture is of a single-layer neural network, when merging and reducing the classes, the within ± 1 2-fold dilution accuracy, the recall, the precision, and the VME reach significant levels. Improvements may be achieved by considering deep neural networks ^5^, however, it is outside the scope of this analysis.

Since we have used *Salmonella* genome data and metadata collected from PATRIC platform, these models are likely useful in the particular region of U.S., however, the methodology is more general and can be used to produce AMR predictors in other countries or regions that have a genome database for infection control.

## Acknowledgments

The author thanks Dr. Boris Panes for valuable discussions and Prof. Dr. Nilton Lincopan for manuscript revision.

## Footnotes

1 The raw accuracy is simply the accuracy of measuring the actual MIC.

2 For amoxicillin/clavulanic acid (or co-amoxiclav) there is an implicit MIC ratio of 2/1.

3 These organizations are the CEN/ISO (European Committee for Standardization/International Organization for Standardization), the DIN (German Institute for Standardization), the EUCAST (European Committee on Antimicrobial Susceptibility Testing) and the CLSI (Clinical and Laboratory Standards Institute).

4 Or whether it is susceptible-dose dependent or nonsusceptible in cases that justify it [16].

5 These are Neural Networks with one or more hidden layers.

